# CBX2 is required during male sex determination to repress female fate at bivalent loci

**DOI:** 10.1101/496984

**Authors:** S. Alexandra Garcia-Moreno, Yi-Tzu Lin, Christopher R. Futtner, Isabella M. Salamone, Blanche Capel, Danielle M. Maatouk

## Abstract

The epigenetic mechanisms that regulate the male vs. female cell-fate decision in the mammalian bipotential gonad are poorly understood. In this study, we developed a quantitative genome-wide profile of H3K4me3 and H3K27me3 in isolated XY and XX gonadal supporting cells before and after sex determination. We show that male and female sex-determining (SD) genes are bivalent before sex determination, providing insight into how the bipotential state of the gonad is established at the epigenetic level. After sex determination, many SD genes of the alternate pathway remain bivalent, possibly contributing to the ability of these cells to transdifferentiate even in adults. It was previously shown that loss of CBX2, the Polycomb group subunit that binds H3K27me3 and mediates silencing, leads to loss of *Sry* and male-to-female sex reversal. The finding that many genes in the Wnt signaling pathway were targeted for H3K27me3-mediated repression in Sertoli cells led us to test whether deletion of *Wnt4* could rescue male development in *Cbx2* mutants. We show that *Sry* expression and testis development were rescued in XY *Cbx2*^*-/-*^;*Wnt4*^*-/-*^ mice. Furthermore, we show that CBX2 directly binds the downstream Wnt signaler *Lef1*, a female-promoting gene that remains bivalent in Sertoli cells. Our results suggest that stabilization of the male fate requires CBX2-mediated repression of bivalent female-determining genes, which would otherwise block *Sry* expression and male development.

**Author Summary:** During development, the bipotential fetal gonad can commit to the male fate and develop as a testis, or to the female fate and develop as an ovary. Mutation of the epigenetic regulator CBX2 leads to ovary development in XY embryos, suggesting a critical role for chromatin remodeling during sex determination. However, the epigenetic mechanisms that regulate the male vs. female cell-fate decision in the mammalian bipotential gonad are poorly understood. In this study, we developed a genome-wide profile of two histone modifications that play critical roles during development: H3K27me3 (repressive) and H3K4me3 (active). We find that sex-determining genes that are initially co-expressed in XX and XY bipotential gonads are bivalent (marked by both H3K4me3 and H3K27me3) prior to sex determination, poised to engage either the male or female fate. Remarkably, after sex determination, repressed genes that promote the alternate fate remain bivalent. We show that stabilization of the male fate requires CBX2-mediated repression of bivalent female-determining genes, which would otherwise block testis development. Our study provides insight into how the bipotential state of the gonad is established at the epigenetic level, and how the male fate is stabilized by repression of the female fate during sex determination.

## Introduction

Gonadal sex determination, the process by which the bipotential fetal gonad initiates development as either a testis or an ovary, is the first critical step in the development of sexually dimorphic internal and external reproductive organs. Sex determination initiates with a binary cell fate decision within a single somatic cell lineage of the gonad, known as the supporting cell lineage [1, 2]. Cells of this lineage are initially held in a bipotential state in the early fetal gonad, poised between male and female fate. In most mammalian species, including mice and humans, expression of the Y-encoded transcription factor *Sex-determining Region Y* (*Sry*) is required to direct the male fate of supporting cells [3-5]. In mice, *Sry* is transiently expressed in XY progenitor cells from ∼E10.5-E12.5, soon after the gonad is first formed [6, 7]. *Sry’s* primary function is to upregulate its downstream target *Sox9* [8]. *Sox9* upregulation and subsequent maintenance through *Fgf9* leads to Sertoli cell differentiation and establishment of the testis pathway [9]. In the absence of a Y chromosome, the canonical Wnt signaling molecules, *Wnt4* and *Rspo1*, become upregulated in XX progenitor cells (Vainio, 1999, Chassot, 2008). The subsequent downstream stabilization of β-catenin [10] together with upregulation of other transcription factors such as *Foxl2* [11], leads to the differentiation of pregranulosa cells in the ovary.

Importantly, upregulation of either Sertoli-or pregranulosa-promoting pathways is accompanied by mutually antagonistic mechanisms, which are critical for repressing the alternate pathway at the time of sex determination [9, 12]. Mapping the up-or down-regulation of each gene in the XX and XY gonad between E11.0 and E12.0, revealed that many genes associated with the female pathway became female-specific by down-regulation in the XY gonad, while a smaller group of male genes became male-specific by down-regulation in the XX gonad [13, 14]. This data established the importance of gene repression in the initiation of male (or female) development.

Interestingly, repression of the alternative fate is actively maintained throughout adulthood. Evidence for this came from a conditional deletion of *Foxl2* in post-natal granulosa cells, which led to their transdifferentiation towards a Sertoli-like state, and reorganization of the ovary into testicular tissue [15]. Conversely, conditional deletion of the XY-determining gene *Dmrt1* in post-natal testes led to transdifferentiation of Sertoli cells into granulosa cells accompanied by ovarian reorganization [16]. This ability to transdifferentiate to the alternate fate highlights both the highly plastic nature of sex determination, and the importance of repressive mechanisms that can stably transmit the initial fate decision long after commitment in fetal life.

In embryonic stem cells and other pluripotent systems, the Polycomb group (PcG) of epigenetic factors have emerged as critical regulators of cell fate specification by both maintaining pluripotency and cell identity through sustained repression of differentiation programs. The PcG proteins assemble into two multi-protein complexes, Polycomb Repressive Complex 1 and 2 (PRC1 and PRC2), which can functionally cooperate to repress target genes [17]. PRC2 establishes H3K27me3 through its catalytic component EZH1/2 [18, 19]. PRC1 can form either canonical (cPRC1) or non-canonical (ncPRC1) sub-complexes depending on the composition of its subunits. While cPRC1 contains a CBX subunit that binds H3K27me3 and promotes chromatin compaction [20], ncPRC1 complexes lack CBX proteins and their recruitment to chromatin is independent of H3K27me3 [21, 22].

The PcG proteins often work alongside the Trithorax group of proteins (TrxG), which have the opposite role of maintaining transcriptional expression through the deposition of H3K4me3 at active promoters [23]. The co-occurrence of an active and a repressive histone modification at the promoter of developmental genes maintains these so-called “bivalent” genes in a lowly expressed state, poised for activation upon reception of a developmental signal [24-26]. Hence, the cooperative function of PcG and TrxG adds a layer of regulation by fine-tuning the timing of gene expression and stabilizing cell fate commitment throughout cell division.

It was previously reported that loss of the cPRC1 subunit CBX2 leads to female development in XY mice and humans, suggesting that cPRC1 is critical for the correct lineage specification of the fetal gonad. Although it was originally proposed that *Cbx2* acts as an activator of the male fate, we considered the possibility that sex reversal in XY Cbx2 mutants is caused by a failure to repress the female pathway, rather than a failure to upregulate the male pathway. To improve our understanding of the epigenetic mechanisms that regulate sex determination, we compared the genome-wide profile of H3K4me3 and H3K27me3 in the supporting cell lineage of the gonad at the bipotential stage (E10.5) and after sex determination has occurred (E13.5). We found that key sex-determining (SD) genes are bivalent prior to sex determination, providing insight into how the bipotential state of gonadal progenitor cells is established. Surprisingly, repressed SD genes remain bivalent after sex-determination, and at least in males, bivalency is retained even in adulthood, possibly contributing to the highly plastic nature of the gonad long after the initial fate commitment. Our finding that the Wnt signaling pathway is a primary target of of PcG repression in males led us to test the possibility that loss of Wnt signaling could rescue male development in *Cbx2* mutants. We show that *Cbx2* is not required for *Sry* expression as previously proposed, as *Sry* expression and testis development were rescued in XY *Cbx2*^*-/-*^;*Wnt4*^*-/-*^ double knockout mice. Furthermore, we show that CBX2 targets the promoter of the downstream Wnt signaler *Lef1* in embryonic and adult testes, further supporting a role for CBX2 as a stabilizer of repression at bivalent genes that would otherwise drive female fate.

## Results

### XY *Cbx2*^*-/-*^ supporting cells fail to repress the female pathway

It was previously shown that loss of the cPRC1 subunit CBX2 leads to complete male-to-female sex reversal in mice and humans, which was proposed to result from a failure to upregulate *Sry* and *Sox9* expression [27-29]. Although it was proposed that CBX2 activates *Sry*, it remained unclear whether CBX2 is a direct mediator of *Sry*, or whether it functions indirectly by repressing an antagonist of *Sry*. In accord with previous results (Katoh-Fukui et al. 2012), we found that there are fewer SOX9-expressing cells in E13.5 XY *Cbx2*^*-/-*^ gonads (Fig 1A-D). Moreover, testis cords that typically characterize this stage of male development (Fig 1B) were completely absent (Fig 1C), and *Cbx2*^*-/-*^ gonads developed as hypoplastic ovaries. Importantly, we found that at E13.5, XY *Cbx2*^*-/-*^ mutant gonads have both FOXL2- and SOX9-expressing cells (Fig 1C and 1D), including some individual cells that express both markers simultaneously (arrows in Fig 1D). This phenotype suggests that sex reversal is due to a failure to repress the female pathway in XY cells, rather than a failure to upregulate *Sry* and *Sox9*.

**Fig 1.**
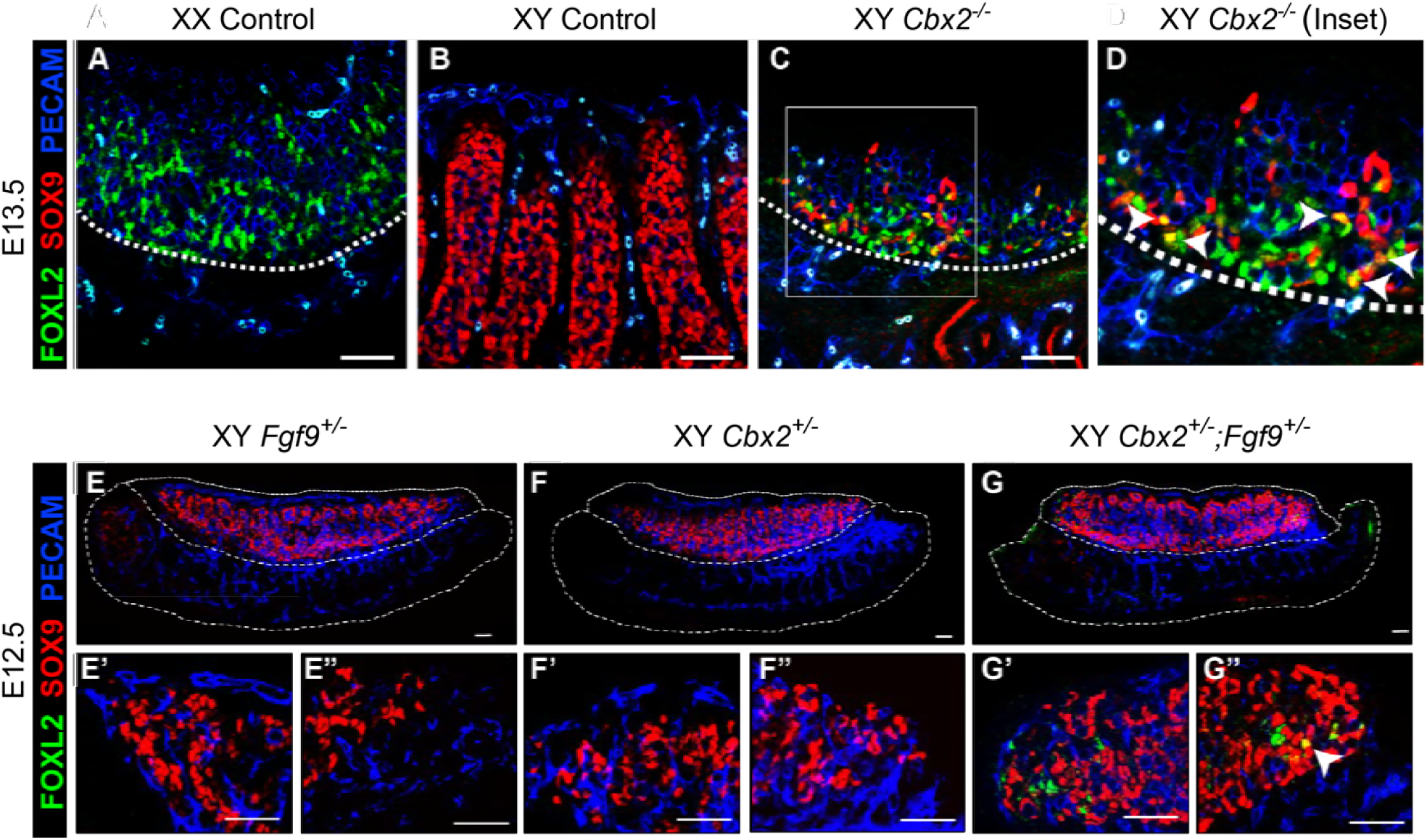
Loss of *Cbx2* in XY cells leads to upregulation of the female pathway. E13.5 (A-D) and E12.5 (E-G”) are stained with the pregranulosa cell marker FOXL2 (green), Sertoli cell marker SOX9 (red), and germ cell and vasculature marker PECAM (blue). WT XX gonads have FOXL2-expressing pregranulosa cells (A), whereas WT XY gonads have SOX9-expressing Sertoli cells (red), which are organized around germ cells forming testis cords (B). (C&D) Loss of *Cbx2* in E13.5 XY gonads leads to reduction of SOX9+ Sertoli cells (red) and gain of FOXL2+ pregranulosa cells (green). Some individual cells express both markers (yellow, arrowheads in D). Testis cords are lost and the morphology resembles WT XX gonads (A). XY gonads of single heterozygotes show no evidence of FOXL2 expression (E, F). The anterior (left, eg. E’) and posterior (right, eg. E”) poles of each gonad are enlarged in the bottom row. Gonads of *Cbx2;Fgf9* double heterozygous mice (G) have FOXL2+ pregranulosa cells at the gonadal poles (G’ and G”). Some individual cells express both markers (yellow, arrowhead in G”). Scale bars represent 50um.

This finding is consistent with several previous lines of work that showed that the male and female pathways are mutually antagonistic. For example, activation of Wnt signaling in the XY gonad can suppress male development [10]. In males, Wnt signaling is repressed (at least partly) by the male-determining gene *Fgf9* [9, 12]. To investigate whether CBX2 synergizes with FGF9 to repress the female pathway, we crossed *Fgf9* and *Cbx2* mutants. Although homozygous loss of either *Cbx2* or *Fgf9* leads to complete male to female sex reversal [27, 30], gonads with heterozygous loss of either gene develop as wild type gonads (Fig 1E-G”). However, gonads of *Cbx2*^*+/-*^;*Fgf9*^*+/-*^ double heterozygotes show partial male-to-female sex reversal at the gonadal poles (Fig 1G”), similar to other models of partial sex reversal [12, 31]. Furthermore, XY double heterozygotes have cells with simultaneous expression of FOXL2 and SOX9, similar to XY *Cbx2*^*-/-*^ gonads (arrow in Fig 1G”). Taken together, these experiments suggest that *Cbx2* functions as a repressor of the female fate during testis development.

### *In vivo* chromatin profiling of gonadal supporting cells

Transcriptional profiling of supporting cells throughout sex determination revealed that at the earliest stages of *Sry* expression, a large network of male- or female-promoting signaling pathways coexist [13]. At this stage, more granulosa-promoting genes are expressed than Sertoli-promoting genes [14]. Commitment to the female fate depends on continued activation of granulosa-promoting genes with little change to Sertoli-promoting genes. In contrast, initiation of the male pathway requires both upregulation of the male pathway and simultaneous repression of the female pathway [13].

Based on the finding that *Cbx2* mutants do not efficiently silence the female pathway (Fig. 1), and the fact that CBX2 is part of a large epigenetic complex that targets hundreds of genes, we speculated that the PcG proteins have a widespread repressive role during male sex determination to establish H3K27me3 silencing marks at genes associated with the female pathway. The PcG proteins functionally cooperate with TrxG proteins, which have the opposite role of maintaining transcriptional expression through the deposition of H3K4me3 [23-26]. To explore the chromatin landscape for these histone marks in XY and XX cells before and after sex determination, we performed ChIP-seq for H3K27me3, H3K4me3 and for Histone 3 (H3) as a means of normalizing across populations [32]. ChIP-seq was performed in FACS (fluorescence activated cell sorted) purified XY and XX bipotential progenitor cells from E10.5 gonads of *Sf1-GFP* transgenic mice [33], committed Sertoli cells from E13.5 gonads of *Sox9-CFP* transgenic mice [34], and committed pregranulosa cells from gonads of E13.5 *TESm-CFP* transgenic mice [35] (Fig 2A). Two independent replicates were performed (each replicate contained pooled cells from multiple embryos), and further validated by ChIP-qPCR (S1 Fig).

**Fig 2.**
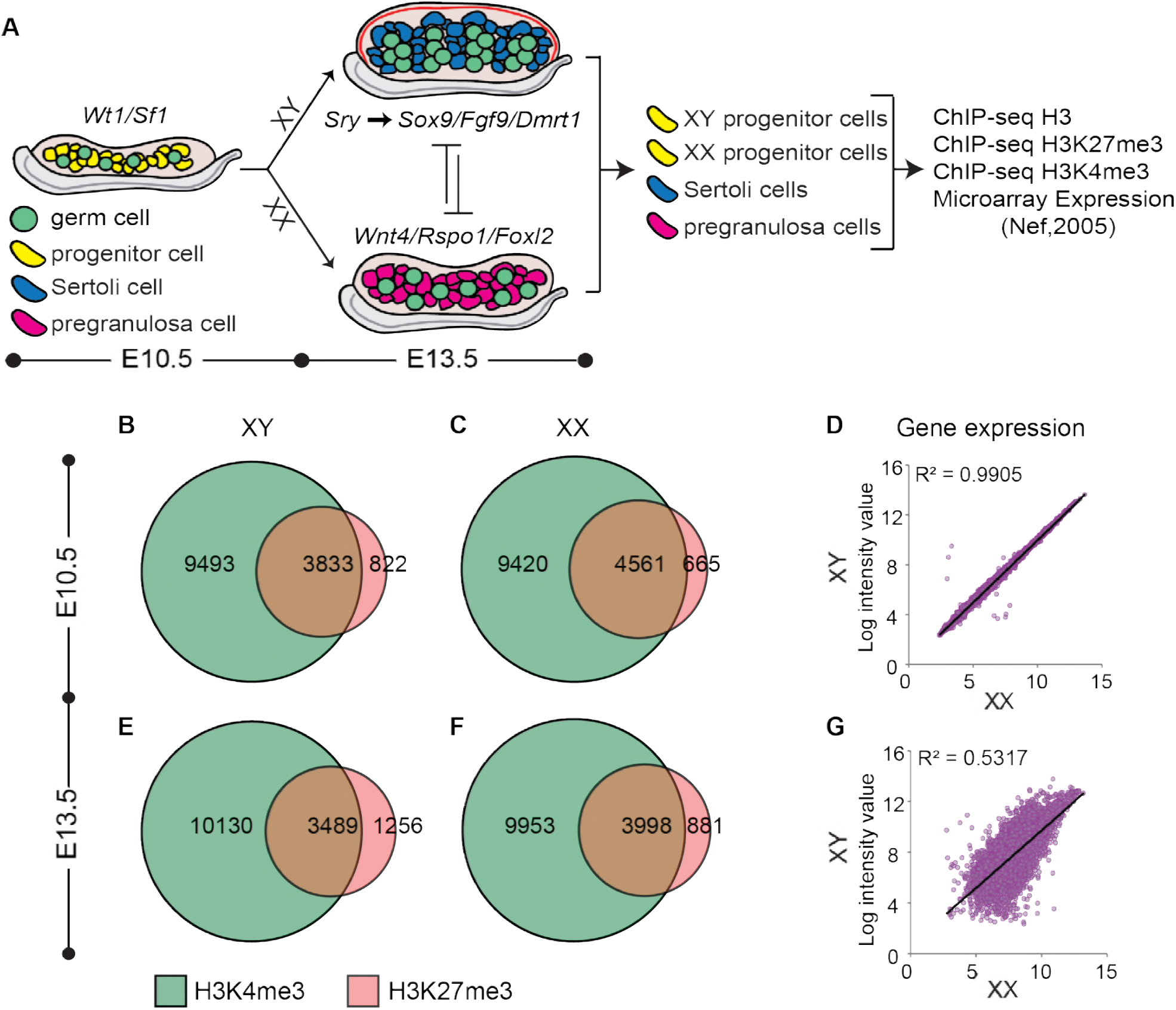
Numbers of bivalent promoters decline as expression becomes sexually dimorphic. (A) An overview of sex determination and work flow. Briefly, supporting progenitor cells (yellow) are bipotential and indistinguishable between XX and XY gonads. Expression of *Sry* directs Sertoli cell differentiation in the testis through upregulation of male-determining genes such as *Sox9, Fgf9* and *Dmrt1*. Absence of *Sry* leads to upregulation of female-determining genes *Wnt4, Rspo1*, and *Foxl2*, directing differentiation in pregranulosa cells in the ovary. XY and XX progenitor cells, Sertoli cells, and pregranulosa cells were FACS-purified and submitted to ChIP-seq for H3, H3K4me3 and H3K27me3. Further analysis made use of microarray expression data from E10.5 and E13.5 purified supporting cells (Nef, S. et al, 2005). (B-C and E-F) Venn diagrams depicting number of promoters marked by H3K27me3 (red) and H3K4me3 (green) in XX and XY supporting cells at E10.5 (B&C) and E13.5 (E&F). (D&G) Linear correlation between expression profiles of XY (y axis) and XX (x axis) progenitor cells at E10.5 (D), and between Sertoli (y axis) and granulosa cells (x axis) at E13.5 (H) (expression profiles from Nef et al., 2005).

ChIP-seq revealed that the largest subset of promoters in all cell types were marked only by H3K4me3 (42-45%), whereas a smaller subset was marked by H3K27me3, the majority of which also overlapped with H3K4me3 (eg. bivalent promoters) (Fig 2B-F and S2A Fig). To investigate the association between these histone modifications and transcriptional activity in the supporting cell lineage, we made use of the microarray gene expression dataset performed by Nef et. al. [36] in FACS purified XY and XX supporting cells from *Sf1-GFP* transgenic mice at E10.5 and E13.5 [33]. As expected, we found that the average transcription level of promoters marked only by H3K4me3 is significantly higher than the transcriptional level from bivalent promoters and promoters marked only by H3K27me3 (S2B Fig). The difference in the average transcriptional level of genes with bivalent promoters versus those marked only by H3K27me3 is of no or low significance, consistent with previous observations that bivalent genes are expressed at low levels and poised for activation.

Previous transcriptional profiling showed that E10.5-E11.0 XY and XX progenitor cells were nearly indistinguishable (Fig 2D), with only a handful of X- or Y-linked genes differentially expressed at this stage [13, 36]. At E10.5, we found that the vast majority of bivalent genes are shared between XY and XX progenitor cells (3545/3788 (XY) and 3545/4561 (XX)) (S2C Fig). The larger number of female-specific bivalent genes is most likely due to the larger number of sequencing reads obtained from the XX-progenitor samples.

By E13.5, differentiation into either Sertoli or pregranulosa cells is accompanied by the upregulation of a number of Sertoli-promoting or pregranulosa-promoting genes (Fig 2G). Consistent with this, the number of bivalent promoters decreased from E10.5 to E13.5 in both XX and XY cells, whereas the number of H3K4me3-only and H3K27me3-only promoters increased (Fig 2F and 2G, and S2A Fig). Comparison of XX and XY samples showed that there is a reduction in the number of overlapping bivalent promoters between XY and XX cells, from 3545 genes at E10.5 to 2963 genes at E13.5 (S2D Fig). These data suggest that different bivalent genes resolve in Sertoli and pregranulosa cells and promote divergence as they differentiate into distinct cell types.

### Key sex-determining genes are poised at the bipotential stage

Our results revealed that 15-17% of transcription start sites (TSS) in both XX and XY supporting cells were co-marked by H3K27me3 and H3K4me3 during the bipotential stage of gonad development (S2B Fig). Bivalent genes play a crucial role in maintaining pluripotency by fine-tuning the timing of gene expression and ensuring the correct lineage commitment [25]. Despite extensive transcriptional profiling of progenitor cells [13, 36], the mechanisms that establish and maintain their bipotential state for ∼24hrs are not well understood. We hypothesized that key sex-determining genes that drive either Sertoli or pregranulosa differentiation are preferentially in the bivalent category at E10.5, held in a poised state for either activation or repression following expression (or absence) of *Sry*. To investigate this hypothesis, we used the set of genes that become either Sertoli-or pregranulosa-specific at E13.5 [14], surveyed H3K4me3 and H3K27me3 deposition at E10.5, and compared the patterns to control genes whose expression is not specifically associated with the Sertoli or pregranulosa pathway (Fig 3A and 3B). Promoters were differentially enriched for H3K4me3 and H3K27me3 consistent with their expression patterns. For example, the promoter of the constitutively expressed *TATA-Box Binding Protein (Tbp)* was marked by high H3K4me3 and lacked H3K27me3 in both XY and XX supporting cells (Fig 3A-E). In contrast, the promoter of the germ-cell specific gene *Oct4*, which is repressed in the supporting cell lineage, is marked by high H3K27me3 and lacks H3K4me3 (Fig 3A-E). Less than 5% of both Sertoli- and pregranulosa-specific gene promoters were marked by H3K27me3-only (Fig 3F). In fact, the vast majority harbored some degree of both H3K4me3 and H3K27me3 enrichment, and approximately 30% of Sertoli- and pregranulosa-specific genes had “high” enrichment (>2.5 log2 enrichment normalized to total H3) of both histone modification marks (Fig 3A, B, F), suggesting a role for bivalent promoters in maintaining the plasticity of supporting progenitor cells to follow either of two fates.

**Fig 3.**
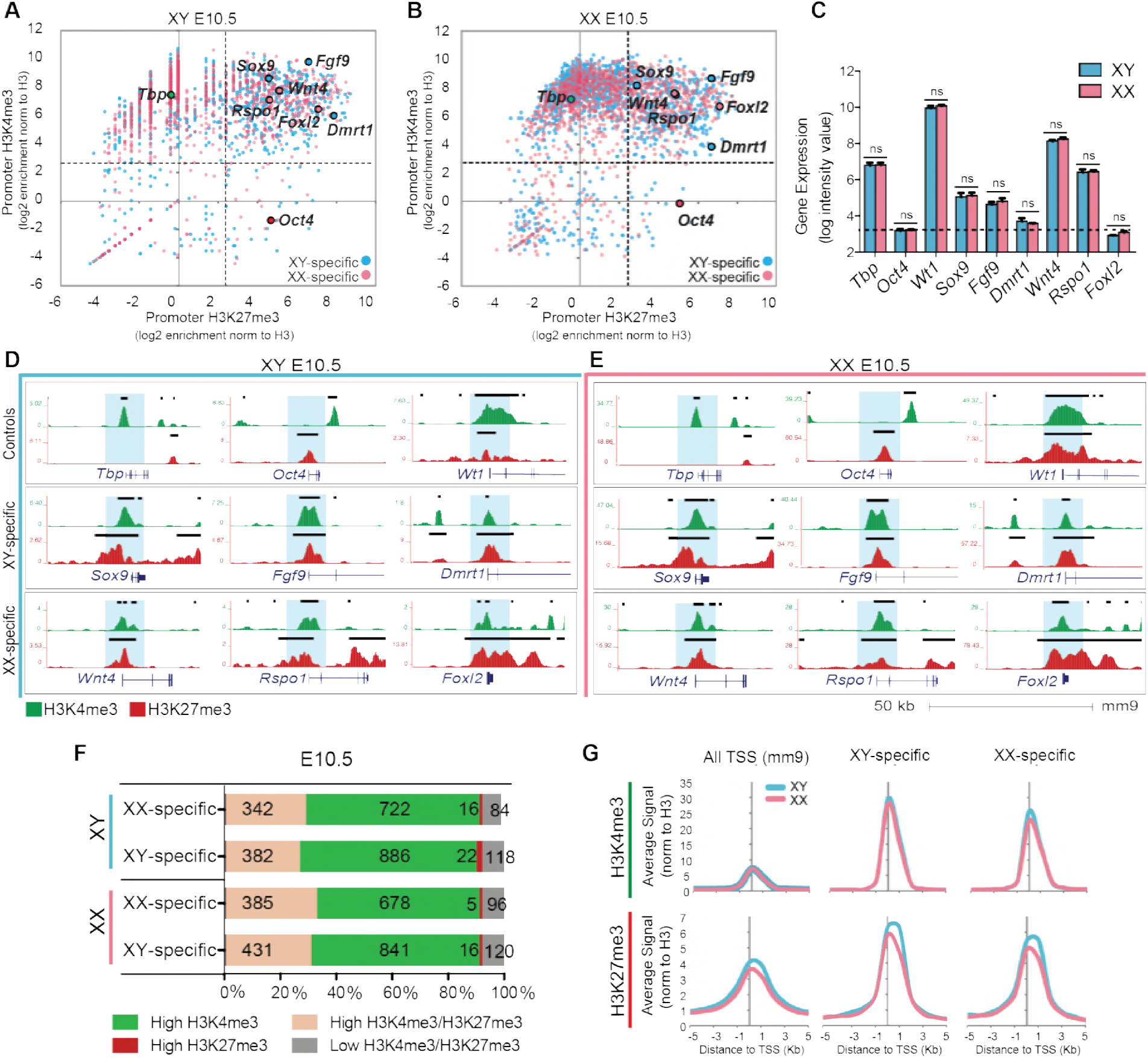
Key SD promoters are marked by both high H3K27me3 and H3K4me3 levels of enrichment prior to sex determination. (A&B) Scatterplots depicting enrichment of H3K27me3 (y axis) and H3K4me3 (x axis) normalized to total H3 within a 2kb promoter region around all TSS (mm9) (grey), in XY (left) and XX (right) progenitor cells. Genes that become either Sertoli- or pregranulosa-specific at E13.5 (Jameson et al., 2012b) are blue or pink respectively. *Oct4* is a germ-cell specific gene that is not expressed in supporting cells; *Tbp* is a constitutively active gene. Note that promoters of many sex determination genes (*Sox9, Fgf9, Dmrt1, Wnt4, Rspo1, Foxl2*) have high (log2>2.5) H3K27me3 and H3K4me3 enrichment. (C) Bar graphs denoting gene expression log intensity values from Nef et al, 2005, for select genes in XY (blue) and XX (pink) progenitor cells. Differences between XY and XX for each gene are not significant as determined by student’s t test. (D&E) Genome browser tracks showing ChIP-seq profiles for H3K4me3 (green) and H3K27me3 (red) in XY (blue, left) and XX (pink, right) progenitor cells. Bold black lines represent significant enrichment when compared to flanking regions as determined by HOMER. Note that SD gene promoters are marked by both histone modifications. (F) Percentage of total promoters marked by high enrichment levels (>2.5 log2 enrichment normalized to H3) of H3K27me3 (red), H3K4me3 (green), both (peach), or neither (grey) throughout sex determination. Numbers of genes in each category are shown (1164 total XX-specific and 1408 total XY-specific genes). (G) Average H3K4me3 (top) and H3K27me3 (bottom) signal normalized to H3 around the TSS (5kb upstream to 5kb downstream) (x axis) at all TSS (mm9), XY-specific promoters, and XX-specific promoters, in XY (blue) and XX (pink) progenitor cells.

In accordance with our hypothesis, promoters of key XY-determining genes such as *Sox9, Fgf9* and *Dmrt1*, and key female-determining genes *Wnt4, Rspo1* and *Foxl2* were bivalent prior to sex determination in both sexes (Fig 3 A-E). Genes such as *Wt1*, which are crucial for gonadal development in both sexe*s*, were also bivalent at E10.5 (Fig 3D and 3E). The observed bivalency in both sexes is consistent with the similar levels of expression between XY and XX progenitor cells (Fig 3C). The enrichment level of H3K27me3 and H3K4me3 modifications of bivalent genes very closely mirrors their gene expression levels. For example, *Foxl2* and *Dmrt1* are repressed at this stage with levels similar to the negative control *Oct4* (Fig 3C), and have a higher H3K27me3/H3K4me3 ratio than other sex-determining genes (Fig 3A and 3B). In contrast, genes such as *Sox9, Fgf9, Wnt4* and *Rspo1* have higher levels of expression (Fig 3C) and, accordingly, a higher H3K4me3/H3K27me3 ratio (Fig 3 A and 3B). However, despite the differences in expression levels, sex determination genes have high enrichment levels of both H3K27me3 and H3K4me3, suggesting that there is transcriptional variability amongst bivalent genes as has been previously reported [24].

With the exception of a few genes linked to the sex chromosomes [13, 14], XY and XX progenitor cells are transcriptionally indistinguishable at E10.5-E11.0 [36]. At this stage, genes that later play a crucial role during sex determination are expressed at similar levels (Fig 3C). Interestingly, in ventral foregut endoderm cells, a population of bipotential cells that give rise to either liver or pancreatic cell types, certain histone modifications can predetermine cell fate [37]. We therefore asked whether gonadal progenitor cells are epigenetically predisposed to their male or female fate. However, no significant differences were observed between XY and XX progenitor cells for mean enrichment levels of H3K4me3 and H3K27me3 at sex-determining genes (Fig 3G), suggesting that promoter deposition of H3K4me3 and H3K27me3 does not direct differentiation of XY and XX progenitor cells, but rather contributes to their bipotential state.

Our results suggest that although only 15-17% of all annotated genes are bivalent at E10.5 (S2A Fig), ∼30% of genes that later become Sertoli-specific as well as 30% of genes that later become pregranulosa-specific are bivalent in both XX and XY progenitor cells, suggesting that bivalency plays a large role in the regulation of sex-determining genes.

### Repressed key sex-determining genes remain poised after sex determination

Having established that key sex-determining genes are bivalent prior to sex determination, we next asked whether their promoters resolve into sex-specific patterns of H3K4me3 and H3K27me3 after sex determination, consistent with the upregulation of either the male or female pathway and repression of the alternate fate. To investigate this, we compared the genome-wide profile of H3K4me3 and H3K27me3 in purified Sertoli and pregranulosa cells from E13.5 XY and XX gonads. We show that upregulation of sex-determining genes is accompanied by a strong reduction of the repressive H3K27me3 mark at their promoters by E13.5 (Fig 4A-F). For example, the key XY-determining genes *Sox9, Fgf9* and *Dmrt1*, which become upregulated in E13.5 XY supporting cells following expression of *Sry* (S3A Fig), show a >2-fold reduction of H3K27me3 enrichment in Sertoli cells (Fig 4F). In total, 260 of the 382 (68%) XY-determining genes that were bivalent at E10.5 lose H3K27me3 enrichment and shift towards an H3K4me3-only state in Sertoli cells, consistent with an upregulation of Sertoli-specific gene transcription (Fig 4A and 4C). Conversely, the key female-determining genes *Wnt4, Rspo1* and *Foxl2*, which become upregulated in E13.5 XX supporting cells in the absence of *Sry* (S3A Fig), show a >2-fold reduction of H3K27me3 enrichment in pregranulosa cells (Fig 4F). Accordingly, 173 of the 385 (45%) female-determining genes that were bivalent at E10.5 shift towards an H3K4me3-only state in pregranulosa cells by E13.5 (Fig 4B and 4C). Promoter H3K4me3 enrichment at these genes does not significantly increase during sex determination, but rather is retained at similar levels (Fig 4F). It is important to note that *Foxl2*, a small single-exon gene, is embedded within an H3K27me3-dense locus and appears to retain H3K27me3 in pregranulosa cells. However, a closer look at the TSS and gene body itself reveals that these regions do in fact lose H3K27me3 upon differentiation (S3B Fig), although this is not evident in our scatterplot analyses that used a 4kb window around the TSS for calculating enrichment levels.

**Fig 4.**
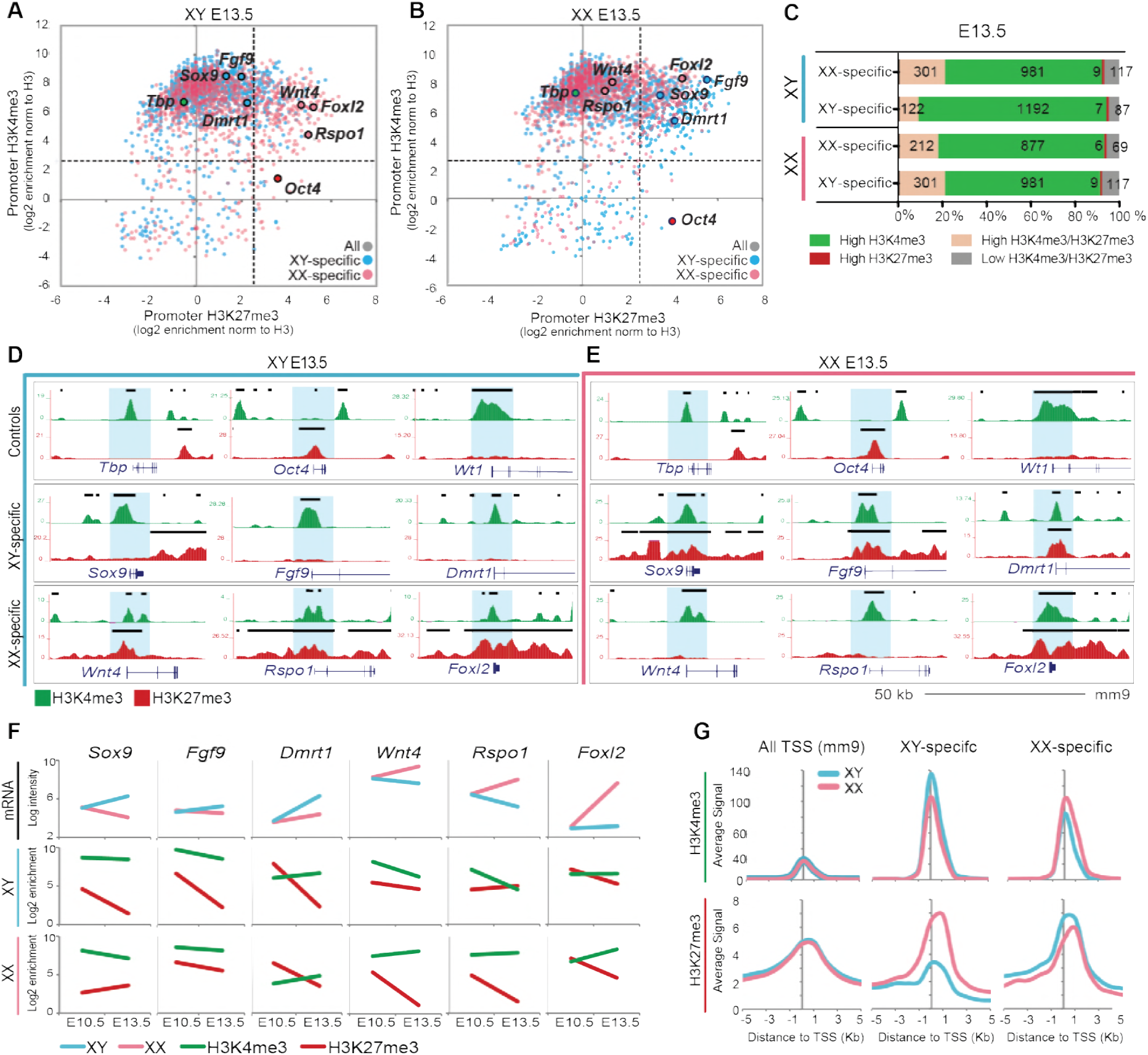
Repressed key SD promoters retain both high H3K27me3 and H3K4me3 levels of enrichment after sex determination. (A&B) Scatterplots depicting enrichment of H3K27me3 (y axis) and H3K4me3 (x axis) normalized to total H3 at a 4kb promoter region in Sertoli cells (left) and pregranulosa cells (right) at E13.5. Genes that become either Sertoli or pregranulosa-specific at E13.5 as determined by Jameson et al., 2012 are in blue and pink respectively. Note that promoters of SD genes that promote the alternate sex have high (log2>2.5) H3K27me3 and H3K4me3 enrichment even after sex determination. (C) Percentage of total promoters marked by high enrichment levels (>2.5 log2 enrichment normalized to H3) of H3K27me3 (red), H3K4me3 (green), both (peach), or neither (grey) throughout sex determination. Numbers of genes in each category are shown (1164 total XX-specific and 1408 total XY-specific genes). (D&E) Genome browser tracks showing ChIP-seq profiles for H3K4me3 (green) and H3K27me3 (red) in Sertoli cells (blue, left) and pregranulosa cells (pink, right) at E13.5. Bold black lines represent significant enrichment when compared to flanking regions as determined by HOMER. Note that repressed key SD genes are marked by both histone modifications. A closer view of *Foxl2* is in S3B Fig. (F) Time course of gene expression (log intensity values from Nef et al., 2005) from E10.5 to E13.5 in XY (blue) and XX (pink) supporting cells (top row), and of H3K4me3 (green) and H3K27me3 (red) promoter enrichment value normalized to H3 in XY (middle row) and XX (bottom row) supporting cells. (G). Average H3K4me3 (top three) and H3K27me3 (bottom three) signal normalized to H3 around the TSS (5kb upstream to 5kb downstream) (x axis) at all TSS (mm9), XY-specific promoters, and XX-specific promoters in Sertoli (blue) and pregranulosa (pink) cells.

Remarkably, we found that sex-determining genes that promote the alternate cell fate and become transcriptionally repressed during sex determination did not lose their active H3K4me3 mark and resolve into H3K27me3-only promoters. Instead, repressed sex-determining genes remained bivalent even after progenitor cells had departed from their bipotential state and differentiated into either Sertoli or pregranulosa cells. For example, *Wnt4, Rspo1* and *Foxl2*, female-determining genes that are actively repressed in Sertoli cells (S3A Fig), retain high enrichment of both H3K4me3 and H3K27me3 (Fig 4A and 4D). Similarly, *Sox9, Fgf9*, and *Dmrt1*, male-determining genes that are not upregulated in pregranulosa cells (3SA Fig), retain high enrichment of both H3K4me3 and H3K27me3 (Fig 4B and 4E). To determine whether this bivalency at repressed genes is retained in adult stages, we performed ChIP-qPCR in purified adult Sertoli cells (> 2 months old males) and found that the female-specific genes *Wnt4, Rspo1, Foxl2, Bmp2* and *Fst* still retained enrichment of both H3K27me3 and H3K4me3 (S4A Fig). Bivalency in adult testes was further validated by ChIP-re-ChIP (S4B Fig).

These results are not exclusive to these key sex-determining genes. In fact, 242 of the 342 (71%) bivalent female-determining genes at E10.5 remain bivalent at E13.5 in Sertoli cells (Fig 3F, and Fig 4A and 4C), and 301 of the 431 (70%) of bivalent male-determining genes at E10.5 remain bivalent at E13.5 in pregranulosa cells (Fig 3F, and Fig 4B and 4C). These results are reflected in the analyses of the mean enrichment level of H3K4me3 and H3K27me3 at the promoters of male- and female-specific genes in Sertoli and pregranulosa cells (Fig 4G). While average H3K4me3 enrichment is higher at male-specific genes in Sertoli cells and at female-specific genes in pregranulosa cells, genes promoting the alternate cell fate have not completely lost H3K4me3, suggesting that even repressed genes retain this active mark. However, the average H3K27me3 enrichment level is higher at female-specific genes in Sertoli cells and at male-specific genes in pregranulosa cells, consistent with its role in maintenance of the repressed state (Fig 4G). Interestingly, there is a significant difference between the mean H3K27me3 enrichment at female-determining genes in Sertoli cells and pregranulosa cells, whereas this difference is not as marked at male-determining genes. This may be a reflection of the total percentage of bivalent genes that have not resolved at E13.5 (Fig 3F and Fig 4C): while 260 of the initial 382 (68%) of XY-specific bivalent genes have resolved in E13.5 Sertoli cells, only 219 of the initial 431 (51%) of XX-specific bivalent genes have resolved in E13.5 pregranulosa cells, possibly because the female developmental pathway is not yet fully established at E13.5 [14]. Interestingly, although levels of H3K27me3 remain constant at genes that retain bivalency from E10.5 to E13.5, the transcription levels of some of these genes decrease (e.g. *Wnt4* and *Rspo1* in Sertoli cells in Fig. 4), pointing towards a possible secondary silencing mechanism.

Our results suggest that key sex-determining genes that promote the alternate cell fate remain in a poised state for activation even after sex determination has occurred, possibly providing supporting cells with an epigenetic memory of their bipotential state and contributing to their ability to transdifferentiate long after sex determination [15, 16].

### Sex reversal is rescued in *Cbx2;Wnt4* double knock-out mice

Our results show that H3K27me3 increases or is stably maintained at promoters of repressed genes during sex determination, with evidence for expansion across loci, a known role of the PcG complex (S5 Fig) [38]. Investigation of pregranulosa-promoting genes in Sertoli cells revealed that genes with known roles in ovary development, such as *Foxl2, Lhx9, Irx3, Lef1, Bmp2, Rspo1, Wnt2b* and *Wnt4*, are amongst the genes with the highest levels of H3K27me3 enrichment (S6 Fig), suggesting that stable repression of these genes is critical in Sertoli cells. Functional annotation of this group of genes (H3K27me3+ pregranulosa genes) revealed that the Wnt signaling pathway is highly represented (S7A Fig), and confirmed that the main targeted biological process is the female reproductive system (S7B Fig). In fact, in addition to *Wnt4* and *Rspo1* (Fig 4), several other members of the Wnt signaling pathway that become female-specific, such as *Wnt2b, Axin2, Lef1*, and *Lgr5* [14], are bivalent with high levels of H3K27me3 at their promoters (S7C Fig). As *Wnt4* is a known ovary-promoting/anti-testis gene [12], our results suggest that the male to female sex reversal observed following loss of *Cbx2* could be substantially due to a failure to repress the Wnt signaling pathway in Sertoli cells.

To test whether loss of *Wnt4* could rescue the male pathway in the *Cbx2*^*-/-*^ XY gonad, we created a *Cbx2*^*-/-*^;*Wnt4*^*-/-*^ double knockout mouse. In accord with our hypothesis, immunofluorescent analysis of E11.5 XY *Cbx2*^*-/-*^;*Wnt4*^*-/-*^ gonads showed that SRY expression was rescued (Fig 5D), similar to wild type XY mice (Fig 5B). Furthermore, SOX9 expression and testis cord formation were rescued in E13.5 *Cbx2*^*-/-*^;*Wnt4*^*-/-*^ gonads (S8 Fig), which develop as testes (Fig 5H and S8D Fig). These results clearly indicate that *Cbx2* is not required for SRY or SOX9 expression as previously proposed. Instead, a failure to repress the female pathway as Sertoli cells differentiate has the indirect effect of blocking the stabilization of *Sry* and *Sox9* expression.

**Fig 5.**
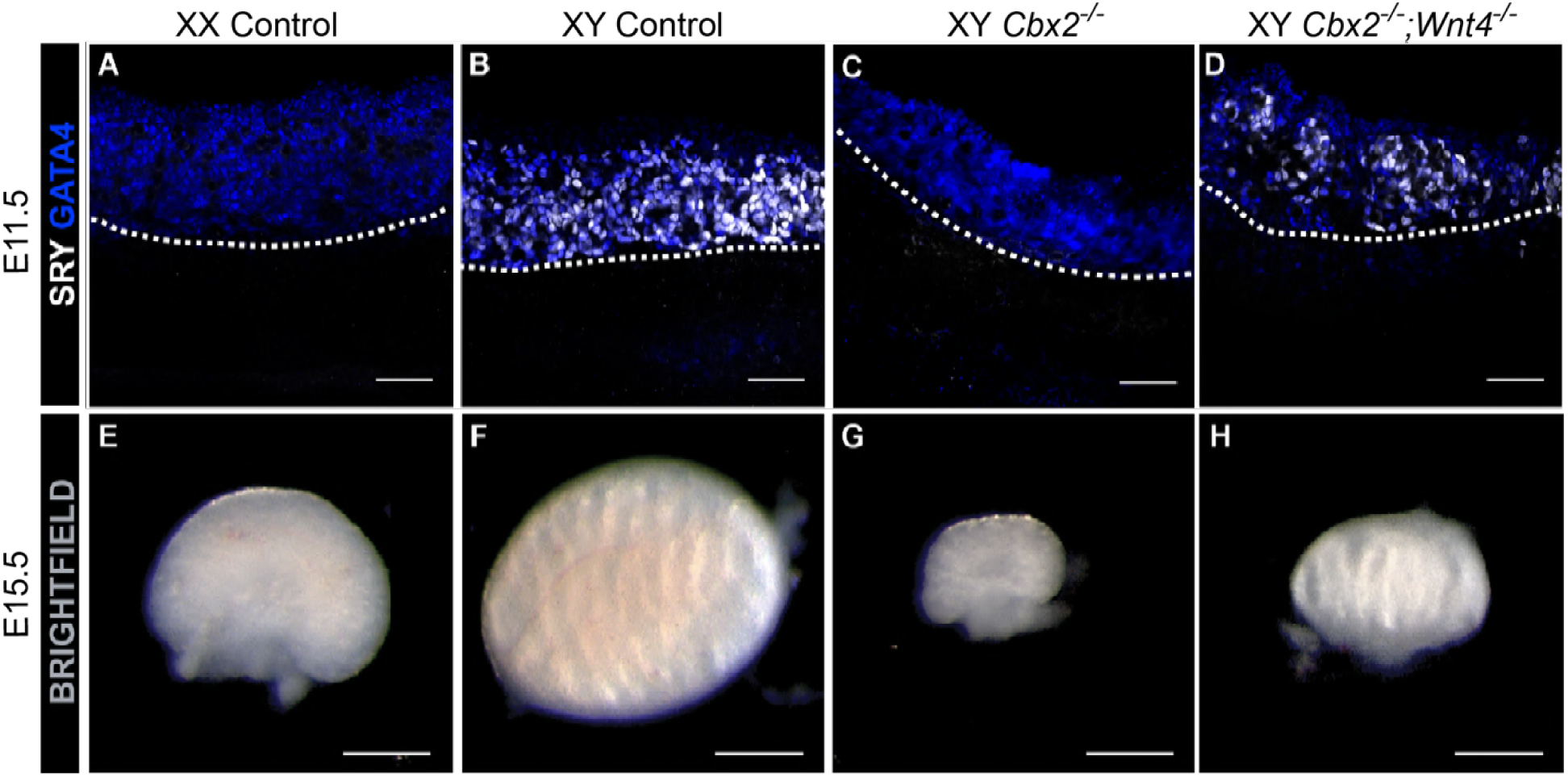
*Sry* expression and testis development are rescued in XY gonads of *Cbx2;Wnt4* double knockout (dKO) mice. Immunofluorescent analysis of gonads from E11.5 (A-D) and brightfield images of E15.5 (E-H) gonads. E11.5 gonads (A-D) are stained with somatic cell marker GATA4 (blue) and SRY (white). Loss of *Cbx2* in E11.5 XY gonads leads to loss of SRY (C). SRY is rescued in *Cbx2;Wnt4* dKO gonads (D). Bright field images of E15.5 gonads show the morphology of WT ovaries (E) and WT testis (F). Loss of *Cbx2* causes male-to-female sex reversal leading to formation of functional but hypoplastic ovaries (G). Loss of *Wnt4* on a *Cbx2* KO background recues testis morphology, but not testis size (H). Images correspond to one of n=3. Scale bars represent 50um (A-D) and 500um (E-H).

It was originally reported that *Cbx2*^*-/-*^ gonads were hypoplastic due to a somatic cell proliferation defect [29]. Despite the fact that XY *Cbx2*^*-/-*^;*Wnt4*^*-/-*^ gonads express testis markers and are morphologically similar to XY wild type gonads (Fig 4D and 4H), gonad size is not rescued, consistent with the idea that *Cbx2* regulates sex determination and gonad size through independent pathways [29]. Our results suggest that CBX2 positively regulates *Sry* and *Sox9* expression by directly repressing the pregranulosa-promoting Wnt signaling pathway.

### CBX2 targets the downstream Wnt signaler *Lef1* for repression

To determine whether CBX2 directly targets *Wnt4* and/or other Wnt signalers, we performed ChIP-qPCR in pooled E13.5 and E14.5 XY gonads (Fig 6A), and in adult (>2m/o) testes (S9 Fig). *Hoxd13*, a known CBX2 target gene [20], was used as a positive control. In contrast, the promoter region of the constitutively active gene *Gapdh* was used as a negative control. Surprisingly, CBX2 did not bind *Wnt4*, or other Wnt signalers such as *Wnt2b, Wnt5a, Rspo1, Axin2, Fzd1*, and *Lgr5*. However, *Lef1* was significantly targeted by CBX2 in both embryonic and adult testes. Downstream of Wnt, LEF1 interacts with β-catenin in the nucleus to drive expression of target genes [39]. During sex determination, *Lef1* is silenced in Sertoli cells and becomes female-specific (Fig 6B) [14, 36]. Accordingly, as *Lef1* is silenced in Sertoli cells, H3K27me3 spreads upstream and downstream off the TSS as well as over the gene body. In granulosa cells, H3K27me3 is removed from the TSS (Fig 6C). Importantly, despite increased levels of H3K27me3, *Lef1* retains H3K4me3 enrichment at its promoter from the progenitor state (Fig 6C). Our results suggest that CBX2 inhibits upregulation of the female pathway during testis development, and possibly maintains Sertoli cell identity in adulthood, by stably repressing WNT4’s downstream target *Lef1*.

**Fig 6.**
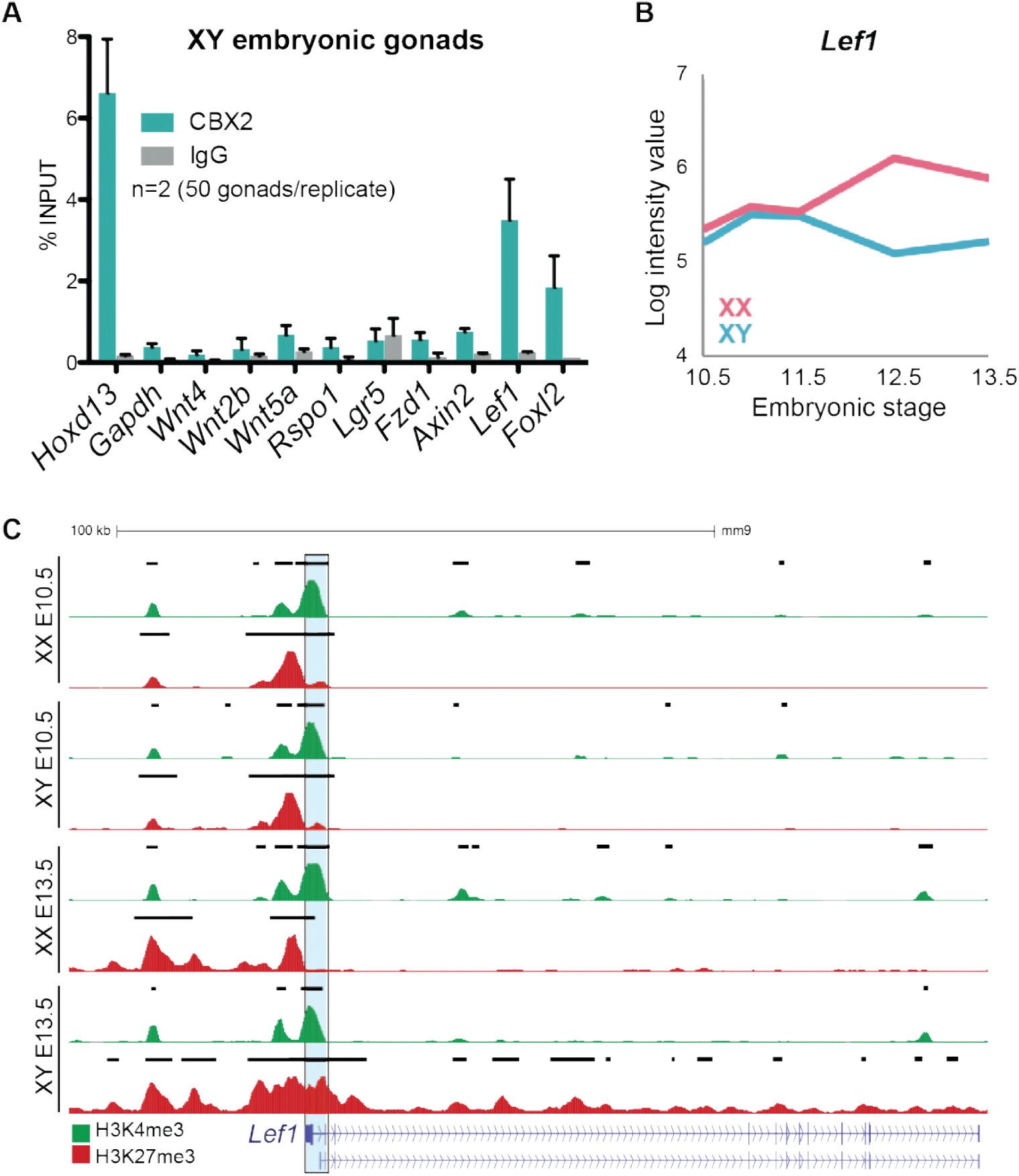
CBX2 targets *Lef1* for repression in XY gonads. (A) ChIP-qPCR for CBX2 on embryonic gonads (2 replicates containing 50 pooled E13.5 and E14.5 XY gonads). * represents p<0.01 as determined by student’s t test when compared to the negative control *Gapdh*. (B) Gene expression profile of *Lef1* in XX (pink) and XY (blue) supporting cells throughout sex determination [36]. (C) Genome browser tracks showing ChIP-seq profiles for H3K4me3 (green) and H3K27me3 (red) at the *Lef1* locus. There are two annotated TSS for *Lef1* in the mm9 genome, however only the 3’-most TSS (blue highlight) loses H3K27me3 in XX E13.5 consistent with its upregulation, suggesting that this TSS is used in the supporting cell lineage. Note the spreading of H3K27me3 over the gene body in XY E13.5. Bold black lines represent significant enrichment when compared to flanking regions as determined by HOMER.

**Fig 6.**
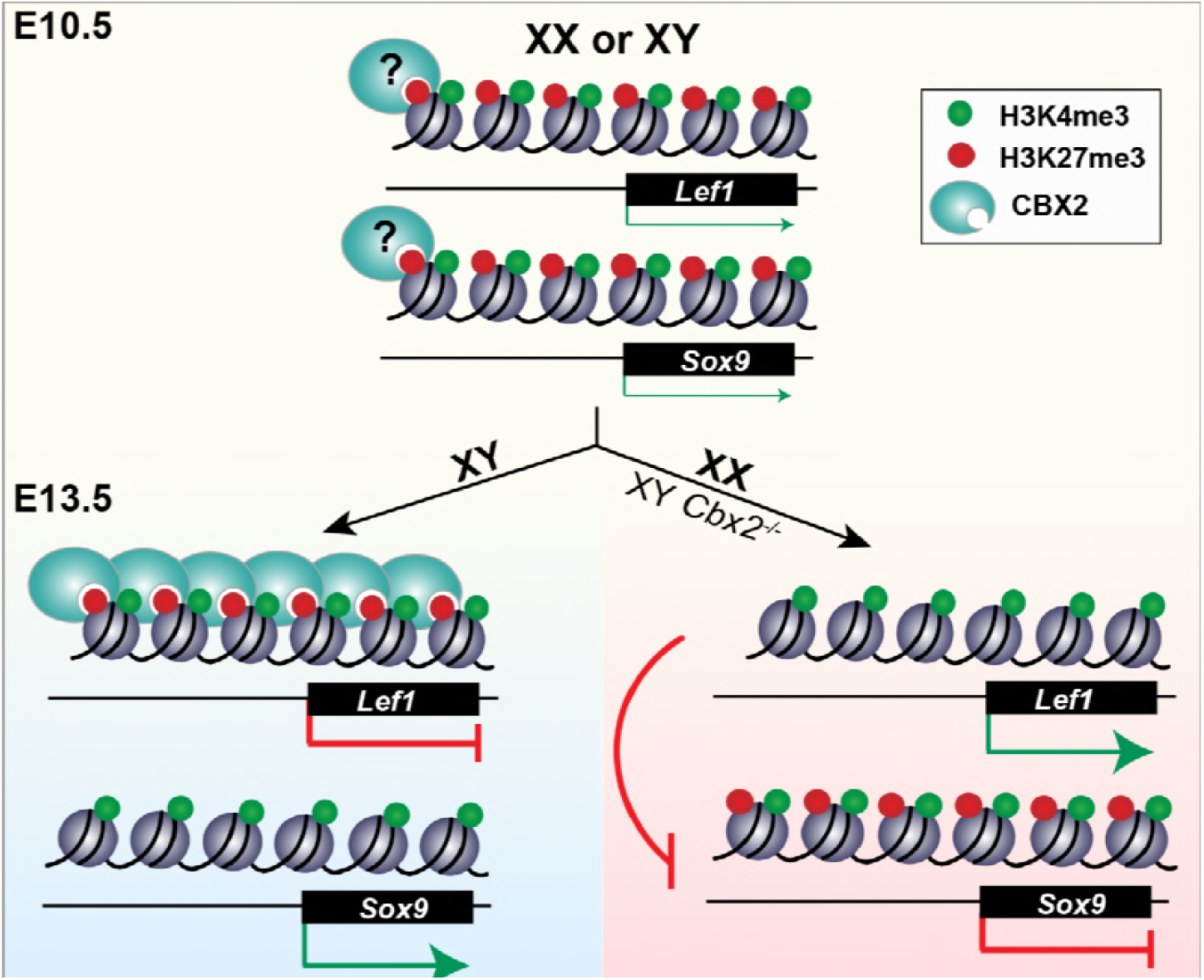
Model of the epigenetic regulation of mammalian sex determination. At the bipotential stage (E10.5), male- (eg. *Sox9)* and female-determining (eg. *Lef1*) genes are bivalent, marked by both H3K27me3 and H3K4me3. Bivalent SD genes are co-expressed at low levels, poised for expression of *Sry* and commitment to the male fate (XY, blue) or in absence of *Sry*, commitment of the female fate through the Wnt signaling pathway (XX, pink). Upregulation of SD genes is accompanied by loss of H3K27me3. Genes that promote the alternate fate and are repressed after sex determination (E13.5) remain bivalent. CBX2 binds to Wnt’s downstream target *Lef1* in XY gonads, inhibiting its upregulation and stabilizing the male fate. In XX E13.5 gonads, or in XY gonads that lack *Cbx2, Lef1* promotes pregranulosa development which blocks upregulation of the male fate (right, pink). It remains unclear whether CBX2 maintains H3K27me3 from the progenitor state in XY cells and is removed from specific targets in XX cells, or whether it is targeted specifically to female genes during Sertoli cell development.

## Discussion

The early progenitors in the bipotential mouse gonad are a valuable model for the investigation of how cells resolve their fate and commit to one of two differentiation pathways. We show that XY cells that lack CBX2, the cPRC1 subunit that binds H3K27me3 and mediates chromatin compaction, fail to repress the female pathway, suggesting that cPRC1 is critical for regulating the male vs. female cell-fate decision of the fetal gonad. To provide insight into the epigenetic mechanisms that regulate this cell-fate decision, we developed an *in vivo* quantitative profile of H3K4me3 and H3K27me3 enrichment in XY and XX gonadal supporting cells at time points before (E10.5) and after (E13.5) sex determination. We show that genes essential to establish both male and female fate are initially poised in both the XX and XY gonad. Once male or female sex determination occurs, the male- or female-expressed bivalent genes resolve into H3K4me3-only promoters. However, the genes associated with the alternate sexual fate retain their bivalent marks, highlighting the importance for a stable repressive mechanism that prevents their upregulation, and ensures that the correct lineage is followed during sex determination. We show that male to female sex-reversal caused by loss of CBX2 can be rescued by simultaneous loss of *Wnt4*, and that CBX2 directly binds *Wnt4’s* downstream target *Lef1*, a female-specific co-factor of β-catenin that remains bivalent in committed Sertoli cells. These findings suggest that *Cbx2* is required during sex determination to stabilize the male fate by blocking the upregulation of bivalent female pathway genes.

The co-occurrence of the active H3K4me3 and the repressive H3K27me3 marks was first reported in ESCs, in which a cohort of developmental genes is “bivalent”, poised for future repression or activation [24-26]. Our findings are consistent with the idea that epigenetic regulation plays a critical role in maintaining pluripotency and facilitating divergence into multiple differentiated states. We show that most genes that become either male- or female-specific after sex determination initially harbor both the H3K4me3 and H3K27me3 modifications at their promoters. More specifically, key sex-determining genes (genes known to cause sex disorders when disrupted) such as the male-determining genes *Sox9, Fgf9* and *Dmrt1*, and the female-determining genes *Wnt4, Rspo1* and *Foxl2*, have high enrichment levels of both chromatin marks at the bipotential stage (>log 2.5 enrichment normalized to total H3). Our findings suggest that sex-determining genes that are initially expressed at similar levels between males and females are bivalent in progenitor cells, held in a chromatin-accessible conformation (Fig 6). This state maintains the balance between the male and female fate yet renders genes responsive to the sex-determining signal triggered by *Sry* in males or *Wnt4/Rspo1* in females.

Typically, upregulation of bivalent genes is accompanied by loss of H3K27me3 at the promoter, whereas repression is accompanied by a loss of H3K4me3 [24]. This binary resolution maintains the transcriptional state throughout cell divisions even in absence of the initial signal. We therefore predicted that sex-determining genes would resolve into sex-specific patterns of histone modifications, stably transmitting the cell fate decision throughout adulthood. Instead, we show that although sex-determining genes that become upregulated do lose their repressive mark, sex-determining genes that promote the alternate pathway and become transcriptionally repressed persist in a poised state even after differentiation into either Sertoli or pregranulosa cells (Fig 6). The finding that female-determining genes retain enrichment of both H3K4me3 and H3K27me3 in Sertoli cells purified from adult testes led us to speculate that transmission of poised sex-determining genes from the bipotential to the differentiated state could confer the remarkable plasticity that enables *in vivo* transdifferentiation of the supporting cell lineage to the alternate cell fate, even long after the fate-decision has been made [15, 16]. Our findings may also explain why repression of the alternate cell fate is an active process throughout adulthood. In the absence of active repressors, poised promoters may be vulnerable to differentiation toward the alternate cell fate. Our data suggest that maintenance of a bivalent but silent state requires the activity of cPRC1.

Accordingly, loss of the cPRC1 subunit CBX2 leads to upregulation of the female pathway and complete male-to-female sex reversal. This was originally thought to be due to a failure to express *Sry*, implying an activating role for PRC1 [29]. Although some PcG proteins have been reported to promote activation of certain targets [40-42], our observation that XY *Cbx2*^*-/-*^ cells co-expressed Sertoli and granulosa markers led us to hypothesize that downregulation of the male pathway is the indirect consequence of failure to repress the female program. Our present results are consistent with this hypothesis and clearly show that 1) *Cbx2* is not required for *Sry* or *Sox9* expression as previously proposed, since *Cbx2;Wnt4* double mutant mice express both of these genes, and 2) CBX2 regulation of *Sry* is indirect through repression of *Wnt4’s* downstream target *Lef1*, which would otherwise promote transcription of downstream pregranulosa-determining genes with anti-*Sry/Sox9* activity (Fig 6).

Multiple genes associated with male development are repressed in XX cells, thus it is still unclear why loss of *Cbx2* does not disrupt ovary development despite its similar expression levels in XY and XX supporting cells [14]. One possibility is that wide-spread epigenetic mechanisms have evolved in males to repress the female pathway, consistent with the fact that more pregranulosa-promoting genes are expressed at the bipotential stage than Sertoli-promoting genes, and that repression of pregranulosa genes is a more significant component of the male pathway [13, 14]. Alternatively, it is possible that maintenance of H3K27me3 at Sertoli-specific genes in XX cells is mediated by CBX subunits other than CBX2, or that cPRC1 achieves specificity by interacting with sex-specific transcription factors or non-coding RNAs during sex determination. Further insight into the mechanistic functions of CBX2 will benefit from more advanced techniques that enable ChIP for non-histone proteins using very small amounts of starting material.

Our results highlight the importance of epigenetic mechanisms in the establishment and resolution of the bipotential state of gonadal precursors. We are the first to develop an *in vivo* genome-wide epigenetic profile of the supporting cell lineage throughout sex determination. We provide insight into how the bipotential state is established, and an explanation for how differentiated supporting cells retain an epigenetic memory of their bipotential state. Furthermore, we describe a widespread role for the PcG proteins in repressing the female pathway during testis development and show that its subunit CBX2 is required to directly repress *Wnt4’s* downstream target *Lef1* in Sertoli cells. Expanding our knowledge of the regulatory mechanisms underlying sex determination may increase our ability to identify the cause of disorders of sex development with unknown etiologies, over 50% of which remain undiagnosed.

## Materials and Methods

(A complete list of primers and antibodies are in the supplemental materials, Tables S1-S4).

### Mice

The *Sox9-CFP, TESm-CFP* and *Sf1-eGFP* reporter mouse lines, and the *Cbx2, Fgf9* and *Wnt4* KO mouse lines were previously generated [30, 34-36, 43] and maintained on a C57BL/6 (B6) background. Timed matings were established between genotypes and CD1 females. The morning of a vaginal plug was considered E0.5. Embryos were collected at E10.5-E13.5 and genotyped by PCR for mutant alleles. Genetic sex was determined using PCR for the presence/absence of a region of the Y chromosome (see primer list for details).

### Immunofluorescence

Embryonic gonads were dissected and fixed for 1hr at RT in 4% paraformaldehyde. Whole gonads were stained as previously described [44] using the following primary antibodies: SRY (1:100, gift of Dagmar Wilhelm, The University of Melbourne, Melbourne, Australia), SOX9 (1:3000, EMD Millipore, Damstadt, Germany), FOXL2 (1:250; Novus Biologicals, Littleton, CO), GATA4 (1:100, Santa Cruz Biotechnology, Santa Cruz, CA), PECAM1 (1:250; BD BioSciences, San Jose, CA). Alexa Fluor488-labeled anti-rat or anti-mouse (Life Technologies, Carlsbad, CA), Alexa Fluor647-labeled anti-goat (Life Technologies, Carlsbad, CA) and Cy3-labeled anti-rabbit (Jackson ImmunoResearch Laboratories, Inc., West Grove, PA) antibodies were used as secondary antibodies (1:500). Nuclei were stained with DAPI. Whole samples were mounted in DABCO (Sigma-Aldrich) in 90% glycerol and imaged with confocal scanning microscopy using a Leica SP2.

### Adult Sertoli cell purification

Sertoli cells were isolated as in [45] using the same solutions and reagents. Briefly, whole testes were dissected from adult B6 males (>2 months old) and tunica was removed. Testis cords were gently separated and rinsed in PBS. Testis cords were then incubated in a collagenase solution to remove interstitial cells. To separate Sertoli cells from germ cells, cords were then incubated in a trypsin solution followed by an enzyme solution containing collagenase and hyaluronidase. At this point, cords were mechanically disassociated by pipetting into a single-cell suspension and washed in PBS. Cells were subjected to a hypotonic shock solution of 1:2 HBSS to H_2_O to lyse germ cells, which are smaller than Sertoli cells. Sertoli cells were recovered by centrifugation at 500rpm for 5 mins at 4°C, washed twice in PBS and resuspended in 500ul of PBS/3% BSA. Cells were then FACS-purified to collect the large cell population while eliminating contaminating germ cells, nuclei and other debris. ∼400K Sertoli cells were routinely recovered from 1 adult mouse. Cells were sorted in PBS/3% BSA for native ChIP-qPCR as described below.

### Chromatin Immunoprecipitation

#### ChIP-seq

Timed matings were set up so that E10.5 and E13.5 XY and XX supporting cells could be FACS-purified and processed for ChIP on the same day. Mint-ChIP was performed with no modifications as in Van Galen *et al*. 2016 on ∼30K–100K FACS-purified supporting cells from pooled gonads. 400K *Drosophila* S2 cells were added per IP as carrier chromatin. Mint-ChIP was performed on two biological replicates. Chromatin was digested using 150 units of MNase (NEB #M0247S) and ChIP was performed using 3μl of H3 antibody (Active Motif #39763), 3μl of H3K4me3 antibody (Active Motif #39159) or 5μl H3K27me3 antibody (CST #9733S).

#### Bioinformatics

Alignment to the mm9 mouse genome was performed using Bowtie. For visualization on the UCSC genome browser, replicates were concatenated to maximize number of reads, peaks were called using MACS2/2.1.0 with the “--broad” setting, and BigWig files were created using bedGraphToBigWig. H3 ChIP was used as input. To identify regions significantly enriched for H3K27me3 compared to flanking regions (peaks), HOMER was used for each independent replicate using the findPeaks function and settings “--style histone”, with a size of 5000 for H3K27me3 and 1000 for H3K4me3. The setting “-C 0” was used in MNase ChIP to disable fold enrichment limit of expected unique tag positions. To limit analyses only to gene promoters, Bedtools Intersect was used to intersect the ChIP-seq called peak files with a file containing all TSSs annotated in the mm9 genome downloaded from the UCSC genome browser, which was expanded to span 1kb upstream and 1kb downstream of the TSS. Tag counts for each 2kb promoter region were obtained using the HOMER annotatePeaks.pl function and were normalized to the sample’s corresponding H3 tag counts. Scatterplots of promoter tag counts normalized to H3 were generated in Excel. Histograms were generated using HOMER’s annotatePeaks.pl function with “-size 20000” and “-hist 1000”. Quantitative comparison was only performed between replicates that were processed together (i.e. between samples in replicate 1 or replicate 2).

#### Native ChIP-qPCR for histone modifications

ChIP-qPCR was modified from the Mint-ChIP protocol described above (Van Galen *et al*. 2016). Briefly, FACS-purified cells or 400K Drosophila S2 cells were resuspended in 20ul 2xPIC/PBS (PIC, Thermo Fisher 1862209) and 20ul 2x Lysis Buffer containing 150u of MNase (NEB #M0247S). Cells were lysed on ice for 20min and chromatin was digested for 10min at 37°C. The digest was stopped by adding 40μl Dilution Buffer + EGTA to a final concentration of 25mM, and incubated on ice 10min. The FACS-purified cells and the S2 cells were pooled, and the sample was brought up to 200μl/IP with Dilution Buffer + PIC. Samples were then split and incubated with 5μl of H3K4me3 antibody (Active Motif #39159), 5μl H3K27me3 antibody (CST #9733S) and 5μl IgG (CST #2729S) as a control overnight at 4°C on a tube rotator. 25μl of protG Dynabeads (Thermo Fisher 10004D) were washed 2x in Dilution Buffer + PIC, added to each IP, and incubated for an additional 3hrs at 4°C on a rotator. Samples were magnetized, and the unbound portion was recovered. Beads were washed 2x in 200μl of ice-cold RIPA, 1x in 200μl ice-cold RIPA/High Salt, 1x in 200μl ice-cold LiCl buffer and 1x in 100μl of TE by moving tubes from the front of the magnet to the back twice. Washed beads were resuspended in 100μl Elution Buffer. Beads and unbound samples were then incubated with 0.5μl of RNase Cocktail (Invitrogen AM2286) and 0.5μl of protK (10mg/ml) at 37°C for 30min and 63°C for 1 hour. DNA was purified using a MinElute Qiagen Kit (28004) and used for qPCR. qPCR was analyzed as Ratio Bound/Unbound=1/2^(Bound CT–Unbound CT), and IgG values were subtracted. Each ChIP-qPCR was performed three times, each on cells from a pool of multiple gonads.

#### Cross-linked ChIP-qPCR for CBX2

(buffers from (Van Galen *et al*. 2016)). Whole adult testes were dissected from >2m/o mice, the tunica was removed, and tubules were dropped into a glass mortar containing 2ml of 1% paraformaldehyde (PFA) /PBS. The timer was set for 10min and testes cords were homogenized with a glass pestle. Homogenized tubules were transferred to a 15ml conical tube and placed on a shaker for the remainder of the 10 minutes. For embryonic gonads, gonads were removed from the mesonephros and directly incubated in 1ml of 1% PFA/PBS on a shaker for 10min at room temperature. PFA was quenched with a final concentration of 125mM glycine and incubated an additional 5min at room temperature. Fixed testes were washed 2x in PBS/PIC by vortexing and centrifuging at 500rpm for 5min at 4°C, and 1x in 1ml PBS/PIC on a rotator for 10min at 4°C. Testes were centrifuged at 500rpm for 5min at 4°C, flash-frozen and stored at −80°C. On day of ChIP, fixed testes were thawed on ice and resuspended in 1ml 1x Lysis Buffer + PIC, homogenized with a plastic pestle that fits a 1.5ml eppendorf tube, and passed through a 35nm cell strainer. Cells were lysed on ice for 30min, and the nuclei were pelleted by centrifugation at 500rpm for 5min at 4°C. The supernatant was removed, and nuclei were resuspended in 300μl RIPA/PIC and incubated on ice 20 min. Chromatin was sonicated to 100-500bp fragments in a Biorupter for 3 rounds of 15min on High, 30s on/off. Debris was pelleted at 14,000rpm for 20min at 4°C. 10% of sheared chromatin was taken as input. 15-20μg of sheared chromatin were incubated overnight with 25μl protG Dynabeads (Thermo Fisher 10004D) pre-incubated for 3hrs with 5μl anti-CBX2 (Bethyl #A302-524A) or anti-IgG (CST #2729S). 50 E13.5 and E14.5 testes were pooled for each ChIP. Beads were washed 2x in 200μl of ice-cold RIPA, 1x in 200ul ice-cold RIPA/High Salt, 1x in 200μl ice-cold LiCl buffer and 1x in 100μl of TE by moving tubes from the front of the magnet to the back twice. Washed beads were resuspended in 100ul Elution Buffer. Beads and input were then incubated with 0.5μl of RNase Cocktail and 0.5ul of protK (10mg/ml) at 37°C for 30min and 63°C for 4 hours to reverse crosslinking. DNA was purified using a MinElute Qiagen Kit (#28004) and used for qPCR. qPCR was analyzed as % Input.

### Ethics Statement

All animal studies were performed under IACUC compliance (protocol # 00003863). Euthanasia was performed by CO2 administration followed by cervical dislocation.

## Acknowledgements

We thank the NU Sequencing core for their help with ChIP-seq; the R. H. Lurie Comprehensive Cancer Center Flow Cytometry Facility at NU, and the Flow Cytometry at the Duke Cancer Center for their help in FACS-sorting our cells; Iordan Batchvarov for mouse maintenance and matings; Tim Reddy and Greg Crawford for sonicator use and Dagmar Wilhelm for the SRY antibody.

## Supplemental Information Legends

### Supplemental Figures 1-9

S1 Fig. Histone marks are consistent between biological replicates and validated by ChIP-qPCR.

S2 Fig. Epigenetic profiling of supporting cells during sex determination.

S3 Fig. Up-regulation of male or female pathway genes is associated with loss of H3K27me3. S4 Fig. Repressed female pathway genes retain bivalent marks in adult Sertoli cells.

S5 Fig. H3K27me3 spreads over repressed loci in Sertoli cells.

S6 Fig. Female-determining genes with critical roles in ovary development are targets of PcG repression in Sertoli cells.

S7 Fig. The Wnt pathway is targeted for H3K27me3-mediated repression in Sertoli cells. S8 Fig. Sex reversal is rescued in *Cbx2;Wnt4* double knockout XY gonads.

S9 Fig. CBX2 targets *Lef1* for repression in adult testes

### Supplemental Table

Complete lists of primers and antibodies.

### Supplemental Excel File

Complete list of Sertoli-specific and pregranulosa-specific H3K27me3+, H3K4me3+ and bivalent genes in XX and XY supporting cells at E10.5 and E13.5.

